# Latent Space Phenotyping: Automatic Image-Based Phenotyping for Treatment Studies

**DOI:** 10.1101/557678

**Authors:** Jordan Ubbens, Mikolaj Cieslak, Przemyslaw Prusinkiewicz, Isobel Parkin, Jana Ebersbach, Ian Stavness

## Abstract

Association mapping studies have enabled researchers to identify candidate loci for many important environmental resistance factors, including agronomically relevant resistance traits in plants. However, traditional genome-by-environment studies such as these require a phenotyping pipeline which is capable of accurately measuring stress responses, typically in an automated high-throughput context using image processing. In this work, we present Latent Space Phenotyping (LSP), a novel phenotyping method which is able to automatically detect and quantify response-to-treatment directly from images. We demonstrate example applications using data from an interspecific cross of the model C_4_ grass *Setaria*, a diversity panel of Sorghum (*S. bicolor*), and the founder panel for a nested association mapping population of Canola (*Brassica napus L*.). Using two synthetically generated image datasets, we then show that LSP is able to successfully recover the simulated resistance QTL in both simple and complex synthetic imagery. We propose LSP as an alternative to traditional image analysis methods for phenotyping, enabling the phenotyping of arbitrary and potentially complex response traits without the need for engineering complicated image processing pipelines.

## 1 Introduction

Developing crop varieties that maintain a consistent yield across different environmental conditions is an important target for plant breeding as weather patterns become more variable due to global changes in climate. Breeding for yield stability requires characterization of an individual plant’s response to biotic and abiotic stress [28] relative to a breeding population. Treatment studies, where individuals in a treated population are subjected to different growing conditions than a control population, play an important role in uncovering the genetic potential for resistance to stress. Such experiments include genotype-by-environment (*G×E*) studies, where the treatment is often abiotic stress, such as water limited growing conditions, or genotype-by-management (*G×M*) studies, where the treatment is different application of inputs, such as herbicide application to assess herbicide tolerance or nitrogen application to assess nitrogen use efficiency. A core challenge for this broad class of experiments, is the ability to quantify and characterize the physical changes observed in the treated plant population relative to the control population, i.e. to phenotype a plant’s *response-to-treatment*.

A number of factors make response-to-treatment a difficult phenotype to quantify. In general, stress affects multiple plant traits simultaneously. Stressors can also have a substantially different type and magnitude of effect on different plant species and different cultivars within the same species. Finally, quantifying response and recovery to stress is sensitive to the timing of observations, and often requires repeated observations over a plant’s life cycle in order to capture important phenological features. An accurate and quantitative assessment of response-to-treatment is particularly important for genomic association studies.

The use of association mapping techniques, such as genome-wide association studies (GWAS) have yielded many candidate loci for agronomically important quantitative traits in plants [4]. For food crops, genome-wide analysis of susceptibility or resistance to abiotic stress factors such as drought [8], nitrogen deficiency [24], salinity [5], or other factors leads to the discovery of genetic differences underlying these agronomically important characteristics. These treatment-based GWAS studies are capable of identifying resistance alleles which could result in a tolerance to a wider variety of environmental conditions if, for example, introgressed into agricultural lines. GWAS studies, however, require large datasets of phenotypic data in order to map associations with genomic data [10].

High-throughput phenotyping (HTP) technologies have advanced rapidly in the past five years to meet the demand for large phenotypic datasets. Recently, image-based HTP has gained popularity, because photographing plants in greenhouses or fields with robots and drones has allowed data collection at yet larger scales. The phenotyping bottleneck has shifted from collecting images, which can now be done routinely, to making sense of those images in order to extract phenotypic information. Although there is a wide selection of software tools available for extracting phenotype information from images [7, 36], the design and implementation of specific phenotyping pipelines is often required for individual studies due to inconsistencies between datasets. This is true of both traditional image analysis where thresholds and parameters need to be adjusted, as well as more recent machine learning techniques which require the time-consuming manual annotation of training data. In addition, some phenotypes are difficult to measure from images, and ad-hoc solutions tailored to a particular imaging modality or dataset are often required in place of more general ones.

To overcome the many challenges associated with image-based phenotyping, we propose Latent Space Phenotyping (LSP), a novel image analysis technique for automatically quantifying response to a treatment from sequences of images in a treatment study. LSP is related to a broad family of techniques known as latent variable models. These models have been previously used for modelling variation in image data via variational inference, using variational autoencoders (VAEs) [14]. LSP instead constructs a latent representation that best discriminates between image sequences of control and treated samples of a plant population and then measures differences among individuals within the latent space to quantify the temporal progression of the effect of the treatment. The key characteristic of LSP in comparison to existing image-based phenotyping methods is that the phenotype estimated using an image analysis pipeline is replaced with an abstract learned concept of the response-to-treatment, inferred automatically from the image data using deep learning techniques. In this way, any visually consistent response can be detected and differentiated, whether that response is a difference in size, shape, colour, or morphology. By abstracting the visual response to the treatment, LSP is able to detect and quantify complex morphological changes and combined changes of multiple phenotypes which would not only be extremely difficult to quantify using an image processing pipeline, but may not even be apparent to a researcher as correlating with the treatment. In this study, we use a combination of natural and simulated datasets to demonstrate that LSP is effective across different plant species (*Setaria*, sorghum, *Brassica napus L*., simulated *Arabidopsis thaliana*) and different types of treatment studies (drought stress, nitrogen deficiency, simulated changes in leaf elevation, and simulated changes in growth rate).

## 2 Materials and Methods

Latent Space Phenotyping consists of a three-stage process as illustrated in Figure 1.1. First, we train an *embedding network* to classify samples as either treated or control based on a sequence of images captured over their growth cycle (Figure 1.1, top panel). During the training process, the embedding network learns image features that best capture how the plants in the dataset respond to the experimental treatment, e.g. drought-stress or nitrogen-deficiency. These embeddings form an n-dimensional latent space, where individual plant images are embedded as abstract n-dimensional points. Second, we train a *decoding network* to perform the reverse process of projecting embedded points from the latent space back to images, in order to obtain a meaningful representation of the latent space (Figure 1.1, middle panel). Finally, we measure response-to-treatment for individual accessions by tracing their path through the latent space from the initial to final time points in their growth cycle. Treated and control replicates of the same accession are expected to have different paths through the latent space. For example, a drought-stressed individual may have a “shorter” path than its control counterpart, or the paths for a treated and control sample may start at similar locations in latent space, but diverge by the final time point in their growth cycle due to visual differences caused by the treatment. Importantly, the embedded paths for control and treatment samples of the same accession are traced and measured in image space (by mapping the path through the decoder) so that these differences are physically meaningful (Figure 1.1, bottom panel). The differences between treated/control samples represent a phenotype for response-to-treatment that is derived directly and automatically from the original image dataset for the experiment. The response-to-treatment phenotype can be used as a trait value with any existing genome-wide association software tool, or interpreted as an objective response rating, e.g. herbicide tolerance, to inform plant breeding decisions. A complete implementation of LSP, called `LSP-Lab`, is provided at https://github.com/p2irc/lsplab.

**Figure 1.1:**
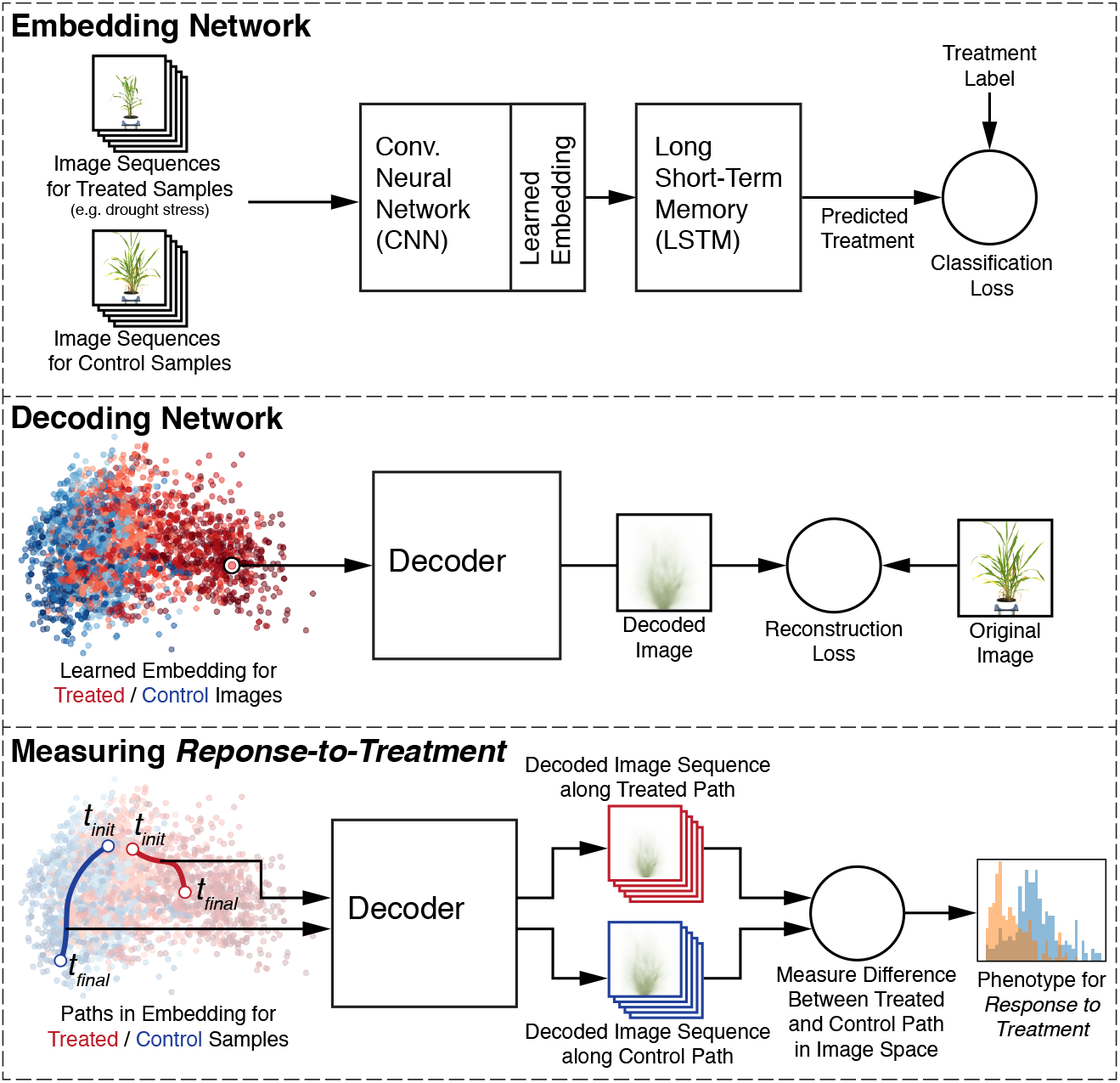
Overview of the proposed technique. The process consists of three phases, which take place in sequence. First, an embedding process projects images into latent space. Second, a decoder is trained to convert these embeddings back into the input space. Lastly, the decoder is used to calculate a geodesic path between the embeddings for the initial and final timepoints.

### 2.1 Dataset Requirements

Performing an LSP analysis requires an image dataset, comprised of images taken at an arbitrary number (*U*) of time points during cultivation for each individual in each of the treatment and control conditions. The initial timepoint should ideally be zero days after stress (DAS), in order to establish this as the baseline for determination of the effect of the treatment. The provided implementation is capable of splitting the analysis into sections of time, for multi-phase experiments. Controlled imaging (using imaging booths, stages, or growth chambers) is recommended in order to maintain consistency in image characteristics such as distance from the camera and the position of the specimen in the frame. However, the method is robust to noise in the images (such as variations in lighting) as long as the noise is consistent between both treatment and control samples, not specific to one condition.

### 2.2 Embedding Network

In order to measure a plant’s response to treatment, it is first necessary to determine which visual characteristics in the images indicate the presence of this effect. To learn the visual features correlating with treatment, LSP utilizes a learned projection of images from a population into an n-dimensional *latent space*, a process known as *embedding*. The embedding is shaped by a supervised learning task, which trains a convolutional neural network (CNN) to extract visual features relevant to the discrimination of treatment and control samples. Performing this embedding allows the method to learn the latent structure of the response, and gives the method the ability to overlook any morphological or temporal characteristics that may be different between accessions, but do not correspond to response to the treatment.

The process of training the embedding network requires only treatment/control labels for each sample. The input to the training process is a sequence of images taken for each individual in the treatment and control conditions. The dataset is divided into training and testing sets with an 80-20 split. Images are standardized by subtracting the mean pixel value and dividing by the standard deviation, and then used as input to a CNN. For each time point image, the activations of the last fully connected layer in this CNN are used as input to a Long-Short Term Memory (LSTM) network (Figure S1).

We describe both CNNs and LSTMs briefly here, but refer to the literature for more detailed summaries of deep learning in general and these network variants in particular [16, 30, 33]. A CNN can be used to learn local feature extractors from image data. The capability of a CNN to learn a complex representation of the data in this way allows the technique to perform well in many complicated image analysis tasks, such as image classification, object detection, semantic and instance segmentation, and many other application areas [16]. CNNs have been used extensively in the recent literature on image-based plant phenotyping, showing promise in several areas, including disease detection and organ counting [12, 21, 22, 25, 33]. For the process of learning an embedding, we implement a simple four-layer convolutional neural network as described in Table S1. Larger architectures were tested and found to show no difference in the experiments reported in this study.

Recurrent neural networks (RNNs) are an extension to neural networks which allows for the use of sequential data. RNNs are a popular tool for time series, video, and natural language problems, for which sequence is an important factor. Briefly, RNNs maintain an internal state which is updated through the sequence, allowing them to incorporate information about the past into the current time point. LSTMs are an extension to RNNs which incorporate a more complicated internal state which is capable of selectively retaining information about the past. LSTMs have also appeared in the plant phenotyping literature, demonstrating that they are able to successfully learn a model of temporal growth dynamics in an accession classification task [30]. LSTMs have also been used as a model of spatial attention in the segmentation of individual leaves from images of rosette plants [27].

The final time point of the LSTM feeds into a two-layer feed-forward neural network, the output of which uses the treatment/control labels of the training images as classification targets, using a standard sigmoid cross-entropy loss for training. The loss on the test set is monitored during training to detect whether or not the embedding network has learned a general concept of the response to the treatment, as opposed to simply overfitting the training data. For the purposes of our application, we prefer embeddings which create only the minimum variance in the latent space necessary for performing the supervised classification task. That is, we prefer embeddings for which variance in most dimensions is close to zero. This helps the subsequent phase of training (Section 2.3) to recover differences in the images which correspond to a generalized concept of response-to-treatment, instead of learning features which are specific to one sample or to a group of samples. To incentivize this, we include an additional loss term for the embedding process alongside the cross-entropy loss and *L*_2_ regularization loss, called the variance loss 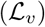,

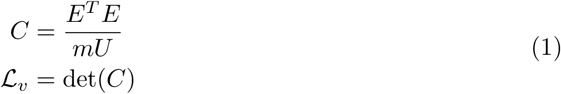

where *E* is the mean-centered matrix of embeddings for a batch of *m* sequences of *U* images. The result is sometimes called the *generalized variance* [35]. In addition, we add a small constant *λ_v_* to the diagonal of *C*. This is for two reasons — first, it prevents the case where zero variance in a dimension causes *C* to be non-invertible, stopping training. Secondly, it stops the optimization from shrinking the variance in one dimension to an infinitesimally small value, effectively pushing the determinant to zero regardless of the variance in the other dimensions and allowing the optimization to ignore the 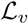 term altogether. Ordinarily, we would find it necessary to restrict the multi-dimensional variance in the latent space by constraining the size of the latent space *n* to the minimum size necessary for convergence. We find that using the variance loss term allows us to use a standard latent space size of *n* = 16 for all experiments, and individual datasets will utilize as few of these available degrees of freedom as necessary as dictated by this term in the loss function. A value of *λ_v_* = 0.2 was used in all experiments. The Adam optimization method [13] is used for training with an initial learning rate of 1e-3.

After the network has finished training, the images of the training and testing sets are then projected into latent space. This embedding is given by the activations of the final fully connected layer in the CNN. In this way, each of the images in each of the treatment and control sequences can be encoded as *n*-dimensional points in the latent space. The final result of the embedding step is all images projected into the same *n*-dimensional space, which can be visualized using a dimensionality reduction technique (e.g., PCA). The embedding plot is used only for visualization purposes, since distances on the embedding plot do not correspond to semantic distance between samples, an issue discussed in Section 2.4. Creating an embedding plot with exact distances between accessions would require calculating on the order of (*Um*)^2^ pair-wise paths in the latent space, which is intractable. However, generating the embedding plot using euclidean distances between embeddings often illustrates stratification of samples in the latent space, albeit with approximate accuracy.

### 2.3 Decoding Network

The second phase of the method involves training a decoder which performs the same function as the embedding process described in Section 2.2, but in reverse. The purpose of the decoder is to define the mapping from latent vectors to image space, discovering the latent structure in the image space, and allowing us to calculate paths in the latent space during the subsequent phase (Section 2.4). The structure of the decoder network consists of a series of convolutional layers followed by transposed convolution layers, which increase the spatial resolution of the input (Table S2). This architecture is similar to those used in other generative tasks, with the exception that there is no linear layer before the first convolutional layer, to prevent the decoder from overfitting. Samples in the training set are projected by the finalized embedding CNN into the latent space, and then the decoder projects these latent space vectors back into the input space (Figure S3). A reconstruction loss function quantifies the difference between the original image and its reconstruction provided by the decoder in terms of mean squared error (MSE). Compared to training the embedding network, a lower learning rate of 1e-4 is used for training the decoder.

Since the embeddings are derived from the supervised classification task, the only features which are encoded in the latent representation are those which are correlated with the response to the treatment. For example, in Figure S3 (middle) the induced angle of the synthetic rosette (the plant leans slightly to the left) is not reflected in the decoder’s prediction, since plant angle is not encoded in the latent space due to it not being correlated with the simulated response-to-treatment. The leaf elevation angle, however, does match between the real and the predicted images. More examples of encoded and decoded images are shown in Figure S7. In practice, the decoder’s output for an input with support in the latent space will tend towards the mean of all images which embed to a point near this location. This mean image should be free of the specific characteristics of any particular accession or individual. The use of MSE creates decoded images which appear blurry — this is an expected result, and helps produce smooth interpolations in the image space when calculating paths in the latent space as described in Section 2.4.

### 2.4 Measuring Response-to-Treatment using the Latent Space

In the third and final part of the process, we seek to quantify the change in the decoder’s output as we travel in the latent space over time, with respect to the embeddings of the images at each time point for a given individual. In other words, we are interested in characterizing the *semantic distance* between decoded images at the initial and final timepoints – that is, the distance between these images in terms of stress propagation. This characterization of semantic distance needs to be considered in terms of the *geodesic path*, on the latent space manifold, rather than euclidean distance in the latent space or in the image space. Figure S2 illustrates the difference between the euclidean distance and the geodesic distance in a hypothetical latent space for a toy example.

In Section 2.3 we defined a decoder (or a *generator function*) 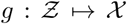 where 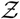 is the latent space and 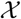 is the input space of the CNN (Figure S3). Since *g* is trivially non-linear, this implies that 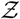 is not a euclidean space, but a (pseudo-) Riemann manifold [2]. The geodesic distance of a path *γ* on a latent space Riemann manifold mapped through a generator *g* in the continuous case is given by

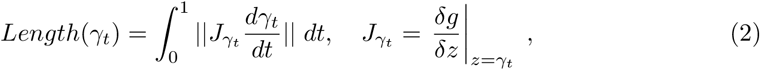

mapping the path through *g* via the Jacobian *J_γt_* [2]. Minimizing this path in the discrete case can be accomplished by optimizing the squared pair-wise distance between a series of intermediate path vertices, minimizing

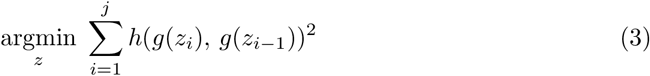

where *h* is a difference function [15]. Performing this optimization on the latent spaces generated by LSP is possible using a standard choice for *h* such as *L*_2_ distance in the image space, since this distance in the image space of the decoder is what we seek to optimize when determining paths through the latent space.

Using the embeddings of the images for the initial and final timepoints provides the start and end points for a path through the latent space. The embeddings of the intermediate timepoints are also computed, and these are used as stationary vertices on the path. Since more vertices means a more accurate discrete approximation of the geodesic path, we interpolate additional intermediate vertices between the stationary vertices. These vertices are calculated by optimizing Equation 3. For all experiments we use as close to, but not more than, 30 vertices for the path, with an equal number of intermediate vertices between each pair of stationary vertices. In general, the choice of the number of vertices is limited by GPU memory. Instead of performing progressive subdivision as in [15], we start from a linear interpolation between stationary vertices. This allows us to perform the optimization all at once, instead of dividing the task into multiple successive optimizations which is potentially more expensive. Calculating the total path length as in the sum in Equation 3 describes the individual in a single unitless scalar value, indicating the difference in semantic distance travelled over the course of the treatment.

Intuitively, the process can be thought of as tracing a path through latent space from where the initial timepoint embeds to where the final time point embeds. In order to find this path, the current position in latent space is decoded into image space by the decoder. Then, the position is moved in the direction which creates the smallest change in this decoded image. As the path is traced in this way, watching the output of the decoder reveals a smooth “animation” where the number of animation frames corresponds to the number of path vertices. The trait value corresponds to how much change there is between each frame and the next, summed up over the entire path. It is important to note that the distance travelled in latent space is irrelevant – the measurement occurs in the output space of the decoder.

When tracing the geodesic path between the first and final timepoints as described, we refer to this as the longitudinal mode. However, it is also possible to perform a crosssectional analysis for populations where there is one treated and one control sample for each accession by tracing a path between the final timepoint for the treated sample and the final timepoint for the control sample. Results for both of the experiments on synthetic data presented here are performed in cross-sectional mode, although longitudinal mode also provides significant results on these datasets. The natural datasets are run in longitudinal mode to match the format of the original study design.

## 3 Results

We evaluated the proposed method using three natural datasets across different plant species and different treatment types, including a population of recombinant inbred lines of Foxtail grass (*Setaria*) treated with drought stress [8], a panel of sorghum treated with nitrogen deficiency [34], and the founders of a nested association mapping population of Canola *Brassica napus* treated with drought stress. We performed two additional experiments using synthetic datasets, including a model of *Arabidopsis thaliana*, where ground-truth candidate loci were verified.

### 3.1 *Setaria* RIL (*S. italica* x *S. viridis*)

We used a published dataset of a recombinant inbred line (RIL) population of the C_4_ model grass *Setaria* [9, 8]. The dataset includes drought and well-watered conditions and has been used to detect QTL relevant to water use efficiency and drought tolerance [26, 8].

The dataset was used as provided by the authors of the original study [8] with a few modifications. The image data was downsampled to 411 by 490 pixels, to allow for a more practical input size for the CNN. Since the camera varies levels of optical zoom over the course of the trial, it is also necessary to reverse the optical zoom by cropping and resizing images to a consistent pixel distance. In order to minimize the effect of perspective shift, the plants were cropped from the top of the pot to the top of the imaging booth, between the left and right scaffolding pieces. This effectively removes the background objects and isolates the plant on a white background. Removing the background is not necessary in the general case – that is, if the background does not change over time. However, since the optical zoom creates differences in background objects, it is practical to remove the background to remove this potential source of noise. The February 1^st^ time point was selected as the initial time point, since many of the earlier time points were taken before emergence. In total, 1,138 individuals representing 189 genotypes and six time points were used. The SNP calls were used as provided by the authors, resulting in a collection of 1,595 SNPs for this experiment. The latent distance values generated by the proposed method were used as trait values for the multiple QTL biparental linkage mapping pipeline provided by the authors of the dataset, in order to replicate the methodology used in the published results.

A histogram of latent distance values for individuals in each of the water-limited and well-watered conditions is shown in Figure 3.1 (left). A total of four QTL were detected with respect to the ratio of the trait under the two conditions. However, we discard these QTL as potentially spurious under the guidance of the original paper [8]. For the difference in trait values between conditions, we identified two QTL associated with drought resistance in the *Setaria* RIL population (Figure 3.1, right). These loci are reported by Feldman et al. as corresponding to plant size and water use efficiency ratio (5@15, within the 95% confidence interval of the reported peak of 5@13.7), and plant size and water use efficiency model fit (7@34), respectively. Although we were only successful in replicating two of the genotype by environment QTL from the published study, many of these previously reported QTL correspond to a water use model incorporating evapotranspiration, not a single trait derived directly from the images such as vegetation area. In order to determine if replications of the same genotype clustered in the output, a one-tailed F test was done on the within-group variance. Of the 378 genotype-treatment pairs, 310 were significantly clustered (*p* < .05).

**Figure 3.1:**
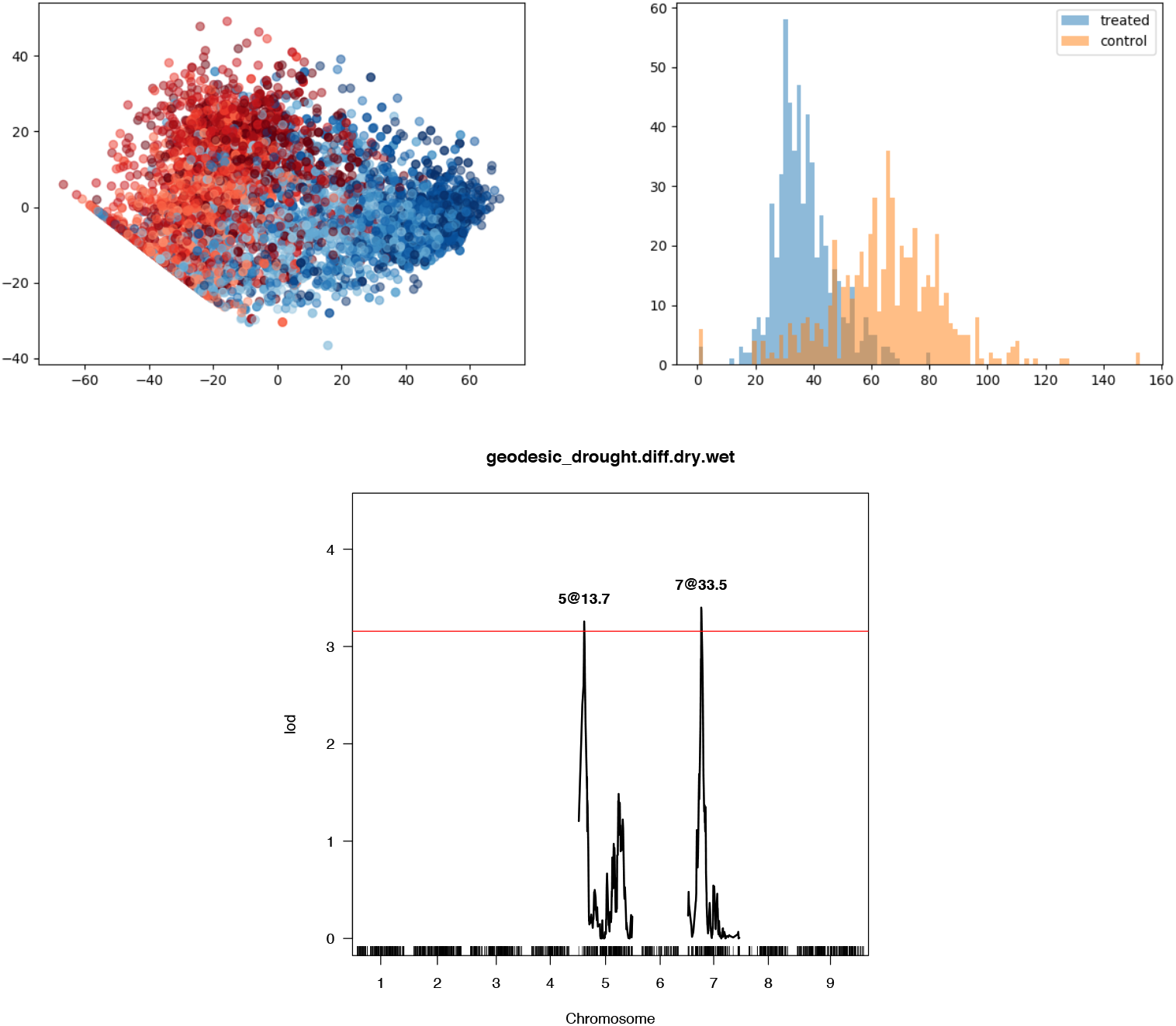
Results for the *Setaria* RIL experiment. Top left: embedding plot. Treated samples are shown in red and control samples are shown in blue. Darker points indicate later timepoints. Top right: histogram of output trait values. Bottom: LOD plot showing QTL for comparison between water-limited and well-watered conditions.

### 3.2 Sorghum (*S. bicolor*)

For this experiment, we used an existing study of nitrogen deficiency in sorghum [34]. The authors of the dataset applied a nitrogen treatment to a panel of 30 different sorghum genotypes. Individuals were placed into the control condition with 100% ammonium and 100% nitrate (100/100), the 50% ammonium and 10% nitrate condition (50/10), or the 10% ammonium and 10% nitrate condition (10/10). Images were analyzed with respect to various shape and color features to detect the presence of a response to the treatment. No association mapping was performed in the published study, and so GWAS results are not provided here. Due to the small size of the dataset, we use data augmentation to help prevent overfitting. This involves introducing random horizontal flips, randomly adjusting brightness and contrast, and cropping to a random area of the image during training. Images were downsampled to 245 by 205 pixels.

In the published study, the authors found that a PCA of 17 different shape features was able to distinguish the control condition (100/100), the high-intensity treatment (10/10), and the low-intensity treatment (50/10). A PCA of various hue and intensity features of segmented vegetation pixels was able to distinguish between the 100/100 and 10/10 treatments, but unable to distinguish between the 50/10 and 10/10 conditions. An LSP analysis of the same dataset was able to distinguish between the 100/100 and 10/10 conditions (Figure 3.2), but failed to converge when tasked with differentiating between the 50/10 and 10/10 treatments. This implies that differences between the two lower nitrogen conditions were too subtle to be detected by either LSP or the collection of pixel intensity features.

**Figure 3.2:**
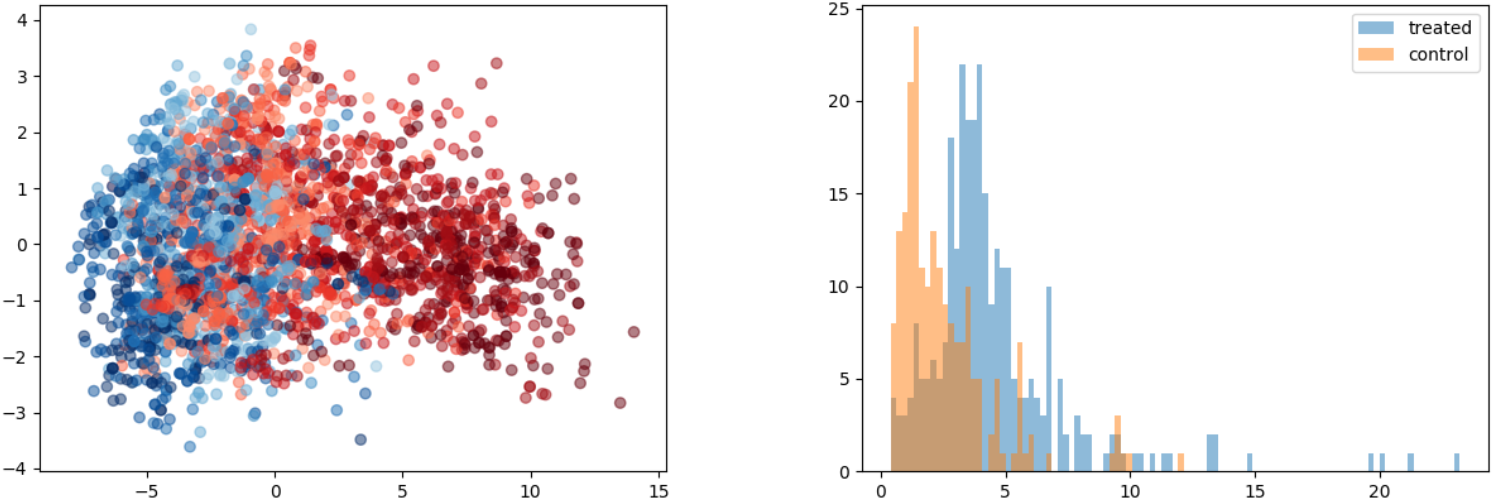
Embedding plot (left) and histogram of generated latent distance values (right) for the sorghum nitrogen treatment dataset for the treated (10/10) and control (100/100) conditions.

### 3.3 Canola (*Brassica napus*)

Next, we performed validation on the founder panel of a nested association mapping population of *B. napus*. In total, 50 genotypes were used in three replications in each of the treated and control conditions, for a total of 300 individuals. Images were taken daily during the early growth period and subsequently every other day, and were downsampled to 245 by 205 pixels. As with the previous datasets, the plants were imaged in a Lemnatec indoor plant imaging system for a total of 40 days and treated individuals were subjected to a drought treatment. In contrast to the *Setaria* RIL experiment, the Canola study involved three phases. First, an initial growth phase which lasted 14 days where no treatment was applied. Next, a 20-day drought phase was applied where watering was reduced from 100% to 40% field capacity, while the pots were imaged every two days. Lastly, a 6-day recovery phase took place where individuals were again watered uniformly across conditions. The results of the LSP analysis are shown in Figure 3.3. Because this experiment involved three distinct treatment phases, the analysis was performed on each of the three relevant portions of the latent space path in series. This gives a separate set of results for each phase, where the performance of individuals can be assessed within each phase. Interpolation was not used on this dataset as it already contains the target number of 30 path vertices.

**Figure 3.3:**
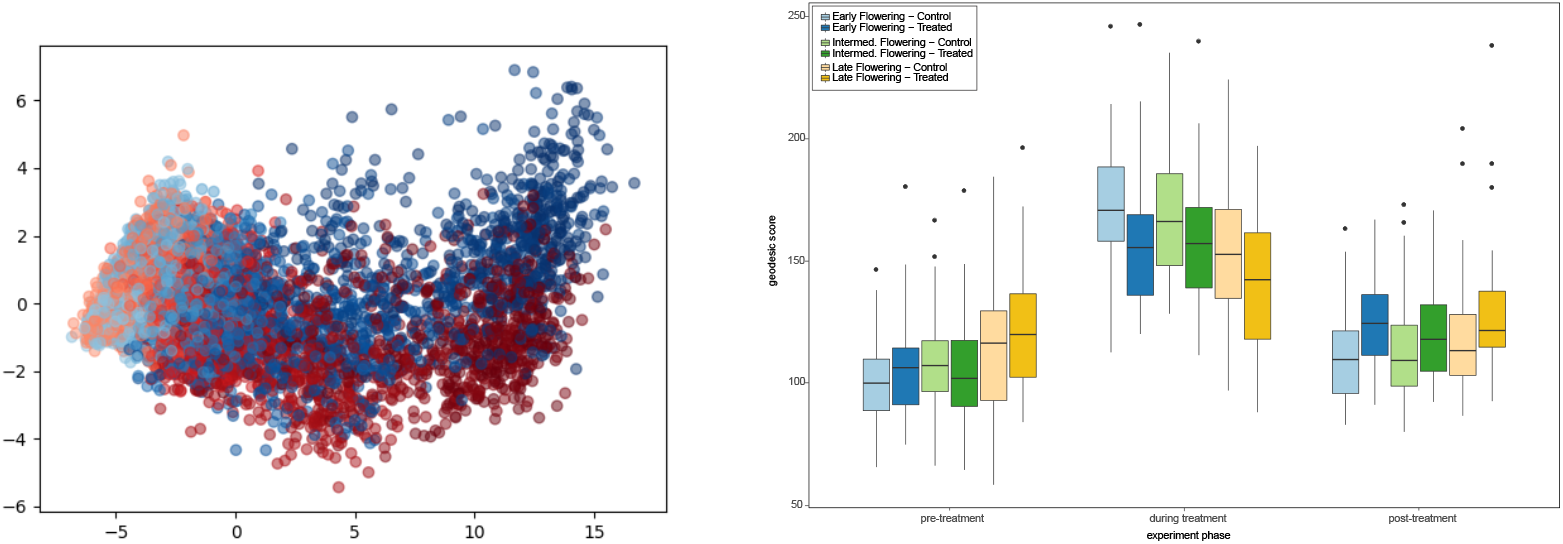
Result for the *B. napus* experiment. Left: embedding plot. Treated samples are shown in red and control samples are shown in blue. Darker points indicate later timepoints. Right: analysis of the output for the three experimental phases, categorized by flowering time.

As with the sorghum trial, the NAM panel of canola is too underpowered to find QTL underlying resistance to drought. However, the trait values output by the proposed method distinguish between the two conditions in the treatment phase. Phenotypic response as measured by geodesic distance was also readily observed when inspecting additional factorial variables, such as flowering time (Figure 3.3, right panel). In order to determine whether multiple replications of the same line were clustered together in the output, a one-tailed F test was performed using the within-group variance for each of the 100 genotype-treatment pairs. For the pre-treatment, treatment, and post-treatment stages, we found that 26, 34, and 29 of the 100 groups were significantly grouped in the output, respectively (*p* < .05).

### 3.4 Synthetic *Arabidopsis thaliana* model

Synthetic images of rosettes [32] and roots [20] have been used previously to train models for phenotyping tasks. Here we used synthetically generated image data as it allows us to introduce specific variance in the imagery based on a simulated casual SNP, and then investigate the method’s ability to recover that variance on the other end by running a GWAS on the simulated population. We use FaST-LMM [19] to perform this analysis and generate the Manhattan plots.

For this purpose, we used an existing L-system-based model of an *A. thaliana* rosette [32], based on observations and measurements of the development of real *A. thaliana* rosettes [23]. The model was run in the lpfg simulation program [1], which simulated the development of the plant over time, and rendered the resulting images. We selected seven of these images corresponding to different time points of the simulation for the LSP analysis.

To generate the synthetic *A. thaliana* genetic dataset, we begin from a real *A. thaliana* genotype database known as the *A. thaliana polymorphism database* [11]. This dataset includes 214,051 SNPs for 1,307 different accessions of *A. thaliana*. A single causal SNP was chosen at random, and we let that SNP represent a polymorphism which confers resistance to a hypothetical treatment that affects the plant’s leaf elevation angle. The elevation angle of the plant’s leaves is sampled from a normal distribution which is parameterized according to whether the sample is untreated, treated-and-resistant, or treated-and-not-resistant. Figure S4 shows the effect of the simulated treatment where the angle of the leaves on the treated plant is increased relative to the untreated sample. Other parameters in the model, such as growth rate, are normally sampled for each accession. It should be noted that, the growth rate of the simulated *A. thaliana* plant is completely uncorrelated from the treatment, as are multiple other model parameters. This means that, although the effect of the treatment is still visually apparent, the embedding network must learn a complex visual concept and cannot rely on measuring the number of plant pixels to discriminate between treated and untreated samples. Since the leaf elevation is modulated as a function of plant maturity, the effect of the treatment is not visible in plants with a low growth rate, adding considerable noise and further increasing the complexity of the task. Also note that performing phenotyping on this image dataset would be challenging, since estimating leaf angle from images is a nontrivial image processing task, especially in the absence of depth information [3, 6].

The method is able to successfully determine the simulated causal locus on chromosome one with no false positives (Figure 3.4). Figure S8 shows a comparison between the proposed method and a naive solution where the image distance is calculated between each pair of images, with no embedding or decoding step. Such an approach would be successful on the simple synthetic imagery described in Section 3.5, but fails in this more complex case.

**Figure 3.4:**
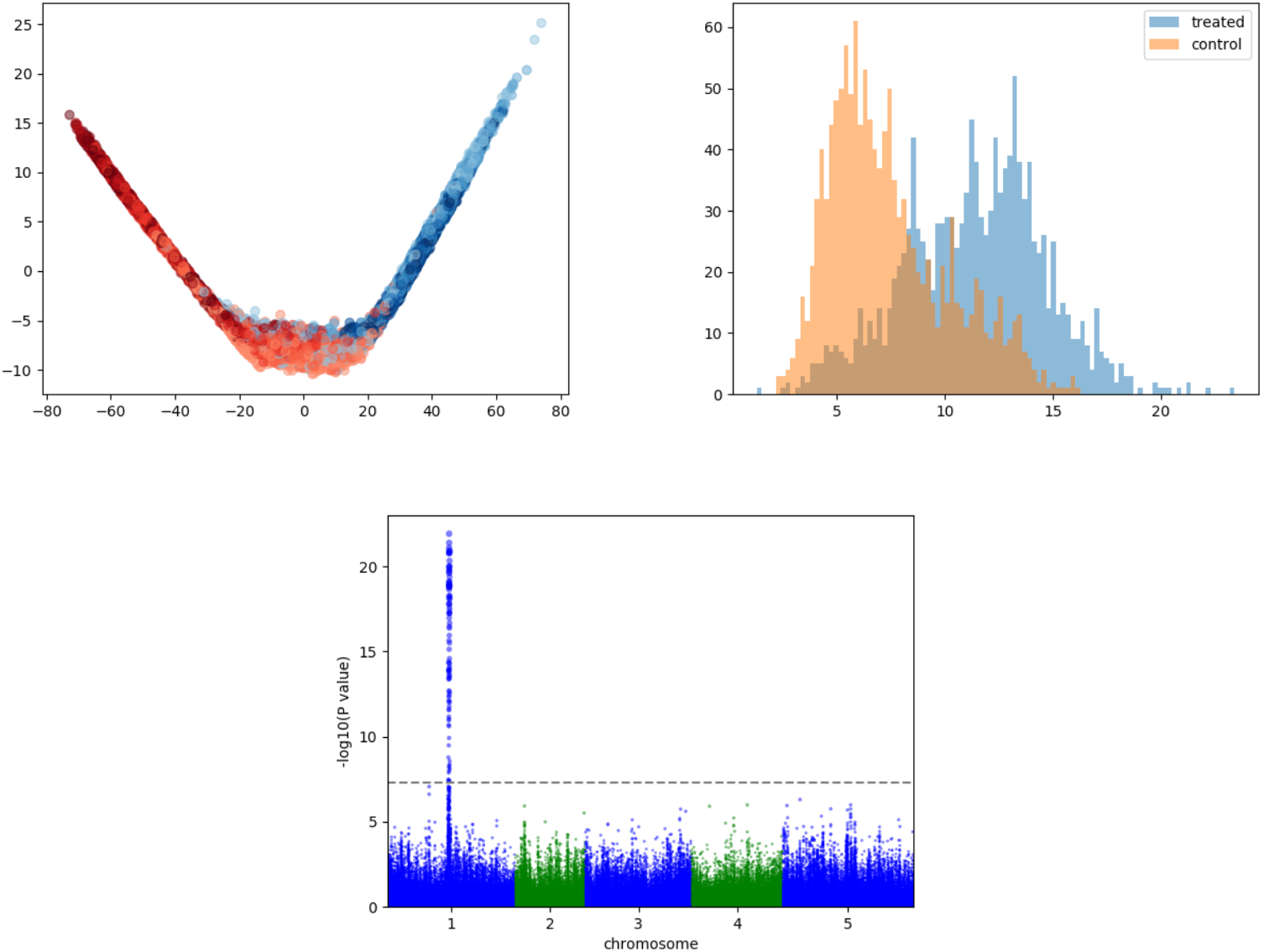
Results for LSP on the synthetic *A. thaliana* model. Top left: embedding plot. Treated samples are shown in red and control samples are shown in blue. Darker points indicate later timepoints. Top right: histogram of output trait values. Bottom: manhattan plot showing the simulated causal locus on chromosome 1. The Bonferroni-corrected *p* < 0.01 significance threshold is shown as a dashed line.

### 3.5 Synthetic circles model

Lastly, we performed an experiment with synthetic imagery intended to show how the LSP method performs under basic conditions, and how its outputs relate to manually-measured phenotypes in this case. For this purpose we use a simple model of the *A. thaliana* rosette which depicts individuals as white circles on black backgrounds, with a hypothetical treatment causing a decreased growth rate of the circle over time in this simple model.

For each of the control and treatment conditions, a sequence of six time points is generated, with images representing a circle growing from an initial diameter (sampled from a normal distribution) to a final diameter. The growth rate of the diameter is drawn from a normal distribution, parameterized according to condition. Additionally, the growth rate under the treated condition is influenced by seven hypothetical resistance loci drawn from a Bernoulli distribution, as well as the presence of the minor allele at a randomly chosen locus in the SNP data. The ground truth growth rate values were used to generate the synthetic imagery.

Performing an LSP analysis of this dataset allows us to forego phenotyping and use the synthetic image data directly. The embedding plot representing the learned embedding of the image data as well as the manhattan plot are shown in Figure 3.5. LSP is able to recover the simulated causal locus with no false positives in this simple application. Relating LSP to the established method of using image processing to extract the growth rate phenotype, we examine the correlation between pair-wise distances in the latent space and differences in measured phenotype between the same accessions. There is significant correlation between calculated geodesic distances in the latent space and the relative growth in the number of white pixels in the synthetic circles dataset (*R* = 0.93, *p* < 0.01).

**Figure 3.5:**
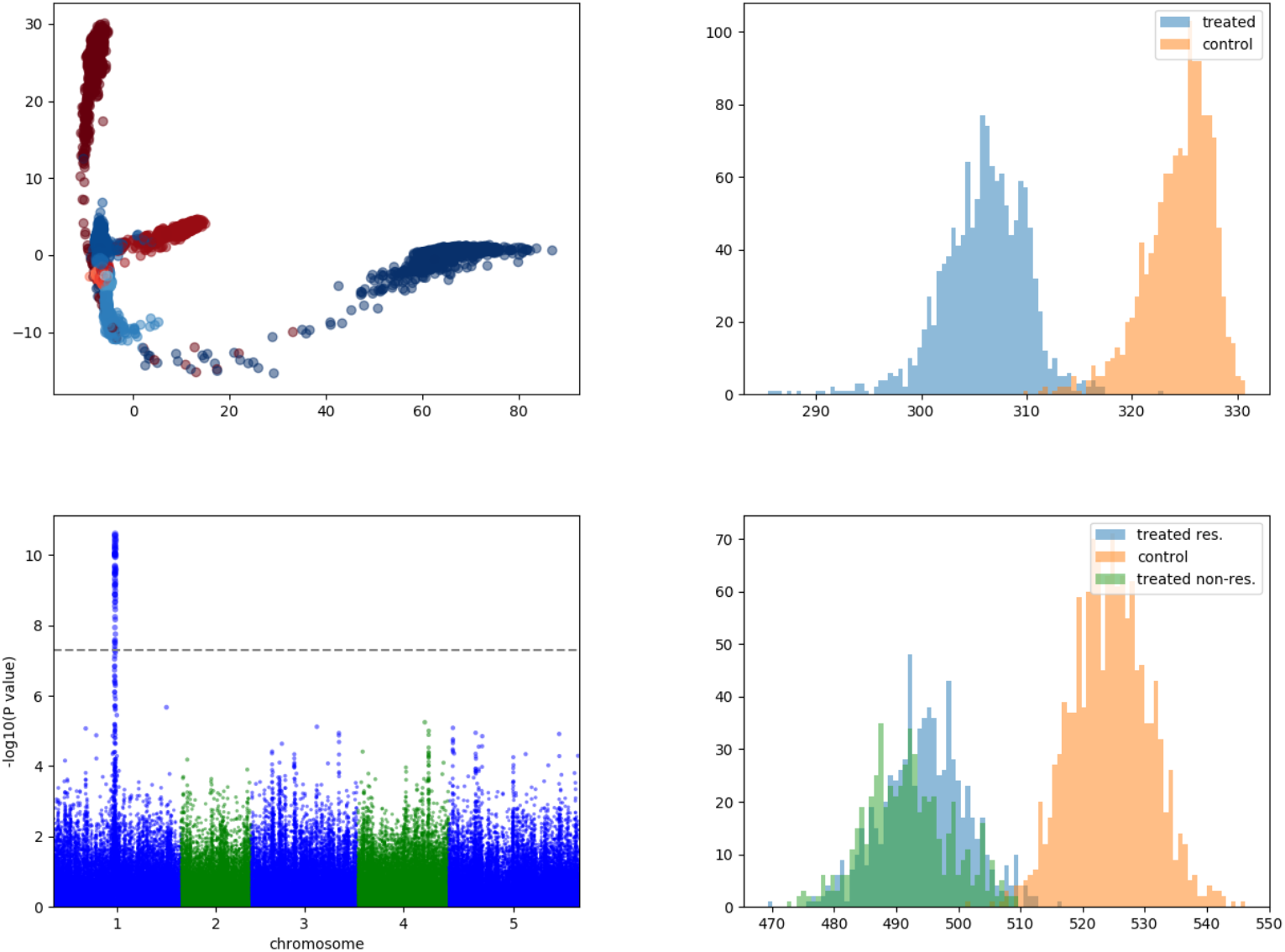
Result of LSP on the synthetic circles dataset. Top left: embedding plot. Treated samples are shown in red and control samples are shown in blue. Darker points indicate later timepoints. Top right: histogram of output trait values. Bottom left: manhattan plot. Bottom right: ground truth growth rate data used to generate the image dataset.

## 4 Discussion

The latent space phenotyping method as described has some limitations, including increased computational requirements compared to the majority of image-based phenotyping techniques. Since the method involves multiple deep neural networks, the use of GPUs is advisable to perform these optimizations in a tractable amount of time. The experiments presented here were performed on two GPUs and the time required per experiment ranged from two to eight hours depending on the number of accessions and the number of timepoints in the dataset. Beyond computational requirements, another limitation of the method is a substantial difference in interpretability compared to GWAS using standard image-based phenotyping techniques. The traits measured with these standard techniques often have a direct and interpretable relationship with the response to the treatment – for example, it has been shown that the number of plant pixels in an image can be used as a proxy for biomass [17]. Therefore, the measured phenotype can be directly interpreted as the biomass of the sample and QTL can be found which correlate with the effect of the treatment on biomass. In the case of LSP, the individual’s response to the treatment is abstracted and quantified only relative to other individuals in the dataset. Interpretability techniques such as saliency maps [29] (Figure S9) can help to elucidate relevant regions in the images, but the measurements still lack a direct biological interpretation in the same way as measurements of biomass. Therefore, candidate loci obtained through LSP must be interpreted differently, and biological explanations must be inferred from the function of the detected loci.

In addition, since the method is non-deterministic due to randomized initial weights and random mini-batching (as with all deep learning methods), repeating the same experiment may output different results. Although there is no guarantee that the trait values reported by the method will be consistent between runs, we found the reported QTL to be consistent across runs for both synthetic datasets. However, a repeat of the *Setaria* RIL experiment resulted in a similar histogram and a between-condition p-value on the same order as the results reported in Figure 3.1, but both previously detected QTL fell below the significance threshold. This is an inevitable consequence of using a non-deterministic method. However, it should be noted that deterministic methods are not inherently repeatable either – different thresholds, outlier detection methods, and data transformations affect the detected loci in these cases.

This work is related to previously described methods which are capable of automatically quantifying differences in morphology between individuals, notably the persistent-homology (PH) method [18]. While PH is focused on automatic shape description, LSP instead learns temporal models of stress which can be dependent on, or independent from, shape. Additionally, PH techniques usually involve the design of a unique system for each shape description task, LSP aims to provide a completely general technique which is not tailored to any particular dataset.

Finally, performing the embedding step can be seen as a type of dimensionality reduction, from the high-dimensional image space to a lower-dimensional embedding space. Doing dimensionality reduction in images has been performed before using techniques such as principal components analysis (PCA) or autoencoders. By performing dimensionality reduction on images, it is possible to recover factors which correspond to pixel variance in the images. For example, the Eigenfaces technique [31] uses PCA on small, greyscale images of faces to learn a series of principle components (PCs). A new image of a face can be encoded as a linear combination of these PCs and this representation can be used to compare the new face against a database of examples to determine similarity. While methods such as Eigenfaces provide a feature vector describing the most major points of variance in the image space, LSP specifically avoids this approach. This is because the most major variations in the images are likely to be from sources completely unrelated to the treatment. For example, the emergence of a new organ creates significant variance in the images, even if the emergence of that organ is not due to the treatment. Using methods such as PCA or autoencoders results in an arbitrary number of features, some or none of which may be useful to the description of the effect of the treatment. Attempting to embed images using other manifold learning techniques such as Multi-Dimensional Scaling (MDS) or Locally Linear Embedding (LLE) suffers from a similar problem – they will attempt to preserve likely meaningless pair-wise relationships in the pixel space.

Let us imagine that one is able to accept the above shortcomings of dimensionality reduction methods such as PCA and autoencoders. Performing the analysis on the full dataset is likely to mostly identify features related to maturity, since this is often the largest cause of variance in the images (and using the full dataset with techniques such as PCA which do not use mini-batching is likely intractable due to memory restrictions). To circumvent this, one could imagine taking the high-dimensional features provided by such methods in separate analyses at each time point. Although most of these features are likely irrelevant, one could use non-parametric significance testing to determine which of the features are correlated with the presence of the treatment. This was validated on both synthetic datasets as well as the *B. napus* dataset using a Mann-Whitney U test. Various PCs were shown to contain information relevant to the condition, and even appeared as describing the most variance (PC 0 and PC 1) in the treatment phase of the *B. napus* trial. However, a more subtle problem with a simplistic latent model and lack of a temporal component is that there cannot be a measurement of the progress of a single, continuous, non-linear process through time, especially if that process contains multiple different stages (such as wilting, followed by senescence). Both PCA and autoencoders are able to encode images and provide reconstructions, making it possible to determine difference in the pixel space given two encoded samples. However, differences in the pixel space can only be calculated between two individuals at discrete time points, and the evolution of these differences across time points cannot be assessed. LSP, on the other hand, integrates stress and maturity in a single continuous space which can be smoothly interpolated through. This space can represent complex, continuous, non-linear changes in different regions. Although PCA was able to detect the presence of the treatment in both the synthetic datasets as well as the *B. napus* experiment, the scores on these significant PCs predictably failed to discriminate between the treatment and control conditions across time.

The results of five experiments demonstrate the capability of LSP to automatically form accurate learned concepts of response-to-treatment from images and recover QTL with a very low false positive rate. As an automated system, the proposed method is exempt from the considerable challenges which arise in developing and deploying image analysis pipelines to first measure phenotypes from images. It is also free from *a priori* assumptions about which visually evident features are caused by the treatment, automatically detecting area, leaf angle, drought stress, and nitrogen stress in five different experiments. Replicating more candidate loci from existing studies will help continue to validate the technique and encourage further study on latent space methods in the biological sciences.

## Acknowledgements

### General

We would like to gratefully acknowledge the Baxter group at the Donald Danforth Plant Science Center for the use of the publicly available *Setaria* RIL and sorghum datasets.

### Author Contributions

J.U. developed the method, performed the experiments, and wrote the manuscript. M.C. and P.P. developed the synthetic *A. thaliana* model and contributed to the manuscript. I.P. developed the *B. napus* dataset and contributed to the manuscript. J.E. performed the flowering time analysis on the *B. napus* experiment and contributed to the manuscript. I.S. supervised the project and contributed to the manuscript.

### Competing Interests

The authors declare that there is no conflict of interest regarding the publication of this article.

### Funding

This research was funded by a Canada First Research Excellence Fund grant from the Natural Sciences and Engineering Research Council of Canada.

### Data Availability

Full datasets and utility scripts needed for reproducing the results and figures presented in Section 3 can be found at https://figshare.com/s/f710381c04c01e2ba319. The data for the *Setaria* RIL experiment and the sorghum experiment are available from the sources referenced by the authors of these datasets [8, 34].

## Supplementary Materials

**Figure S1:**
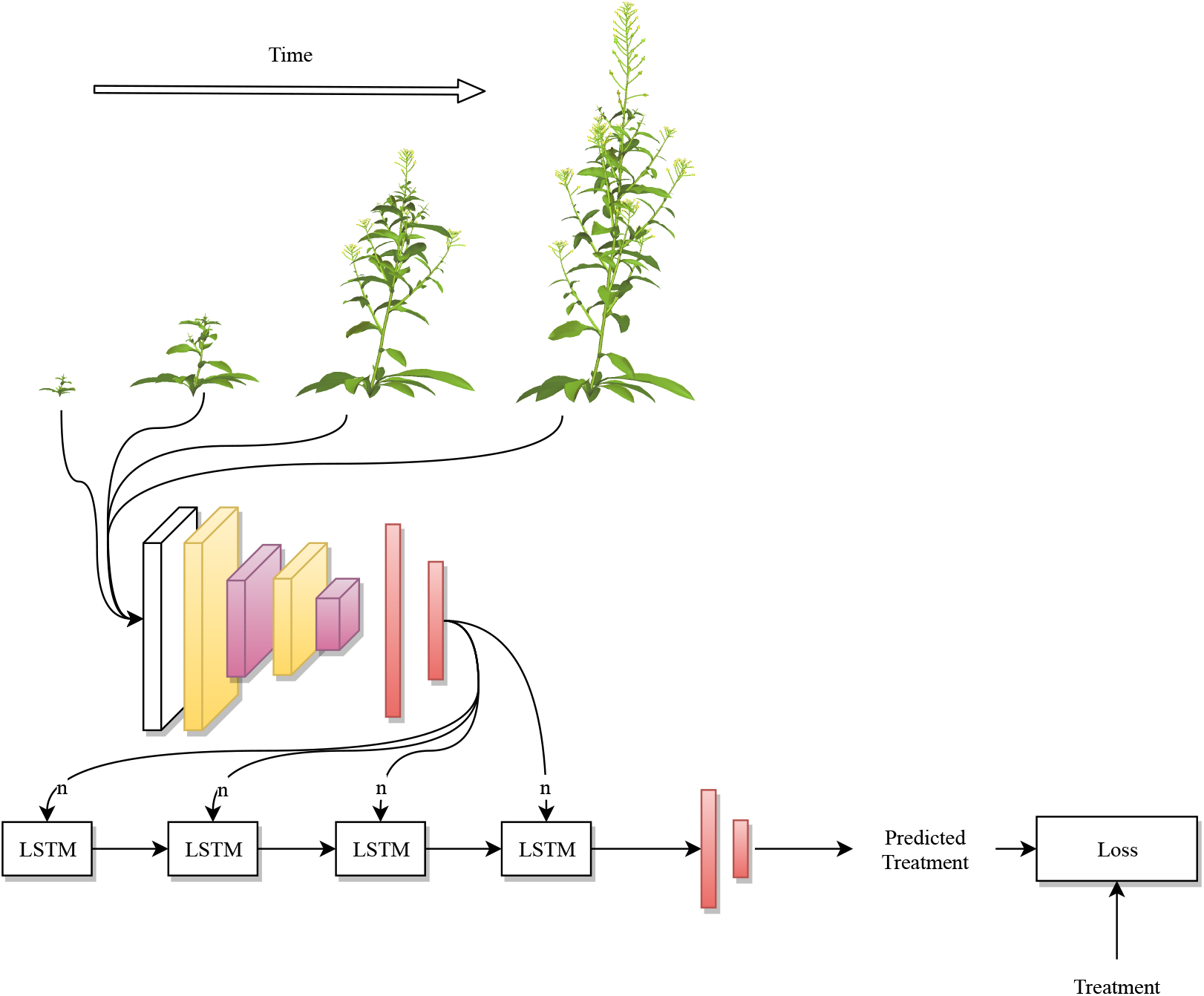
The deep network used in learning an embedding. A CNN takes a sequence of images at various timepoints and feeds outputs to an LSTM, which in turn is used to predict the treatment. The LSTM is removed and the CNN is retained to embed new samples.

**Table S1:**
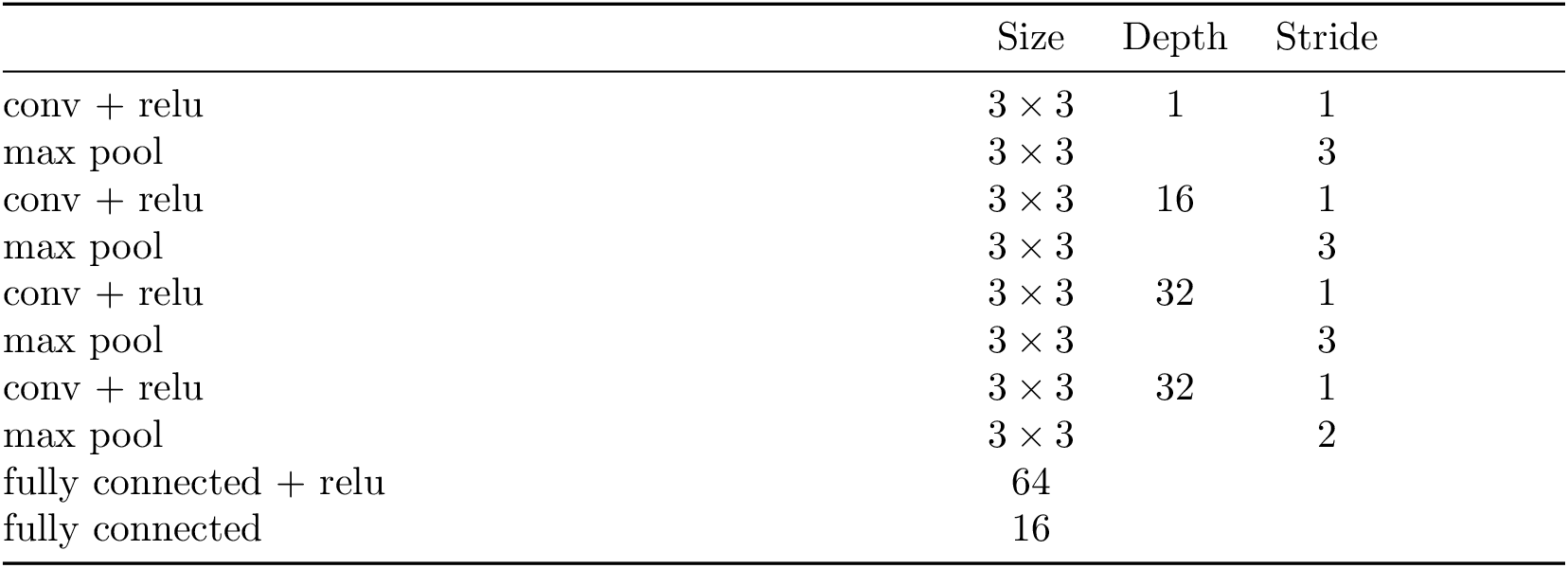
Architecture details for the convolutional neural network used in the embedding. All pooling layers are followed by batch normalization.

**Table S2:**
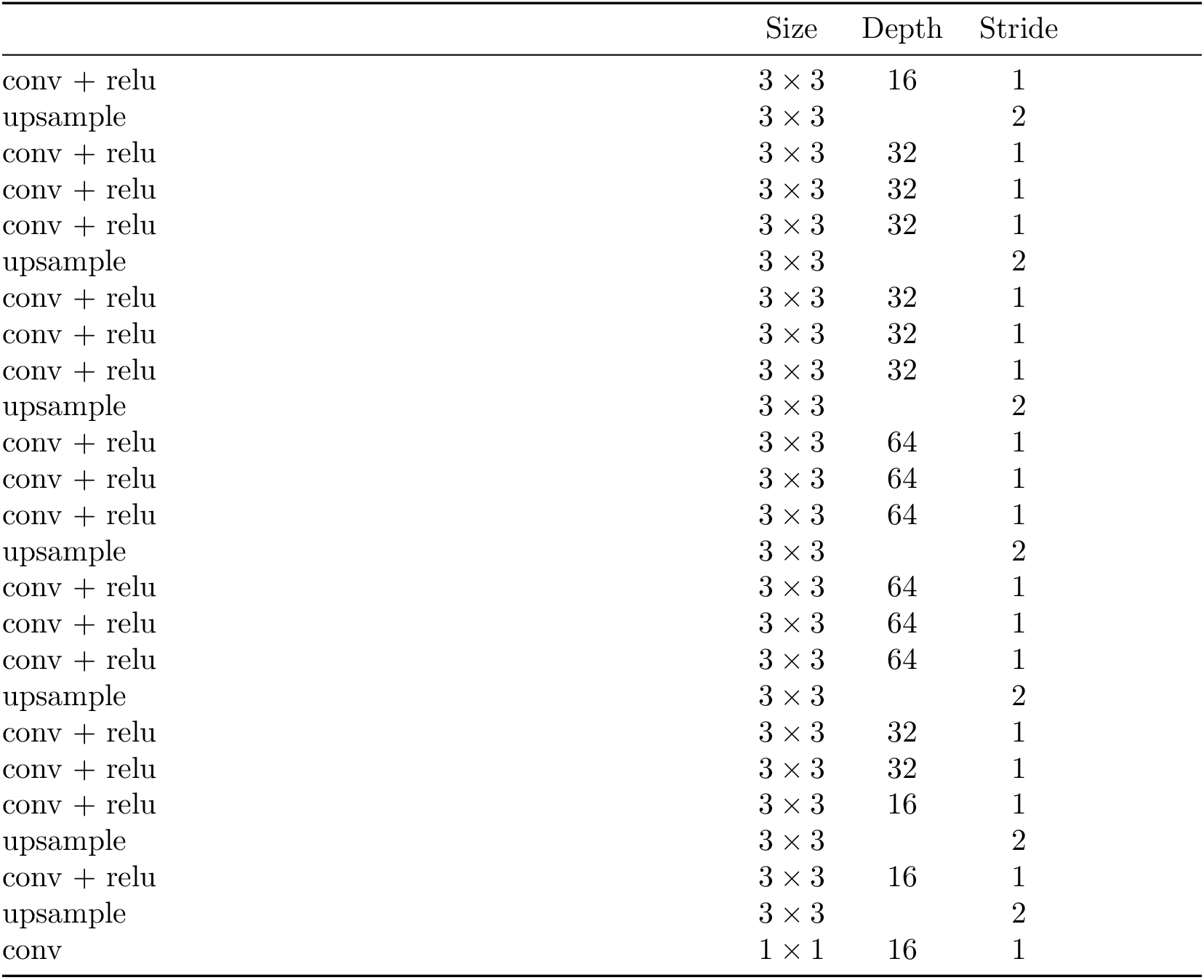
Architecture details for the decoder network. The upsample blocks refer to transposed convolutions.

**Figure S2:**
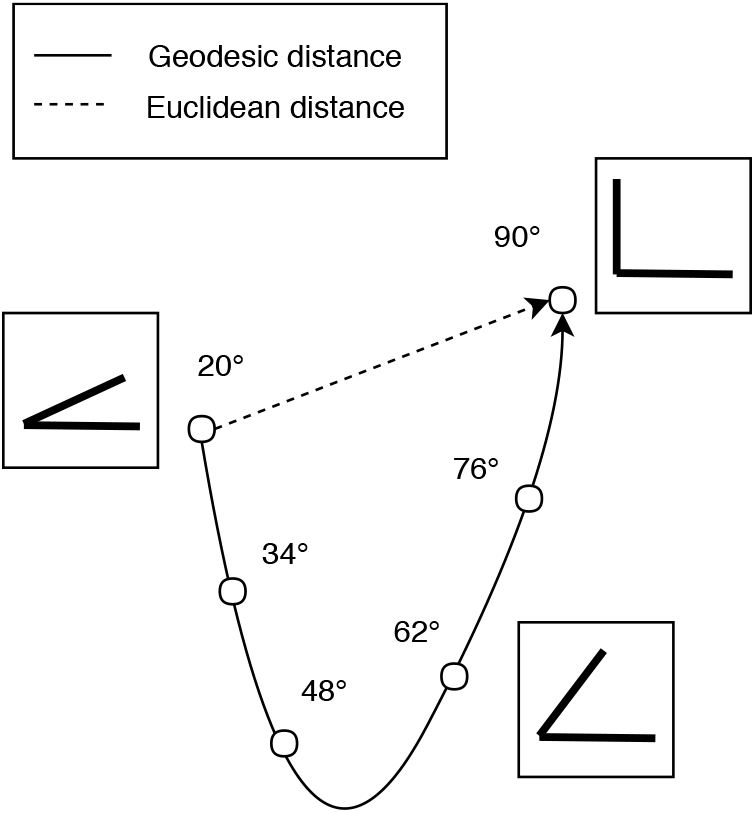
Distance between embeddings of images of lines at different angles in a hypothetical latent space. Here, the *semantic distance* is the difference in the interior angle. The euclidean distance between images in the image space is constant, and the euclidean distance (dotted arrow) between their encodings in the latent space is evidently not representative of the semantic distance. However, the geodesic path (solid arrow) between images represents the semantic distance well.

**Figure S3:**
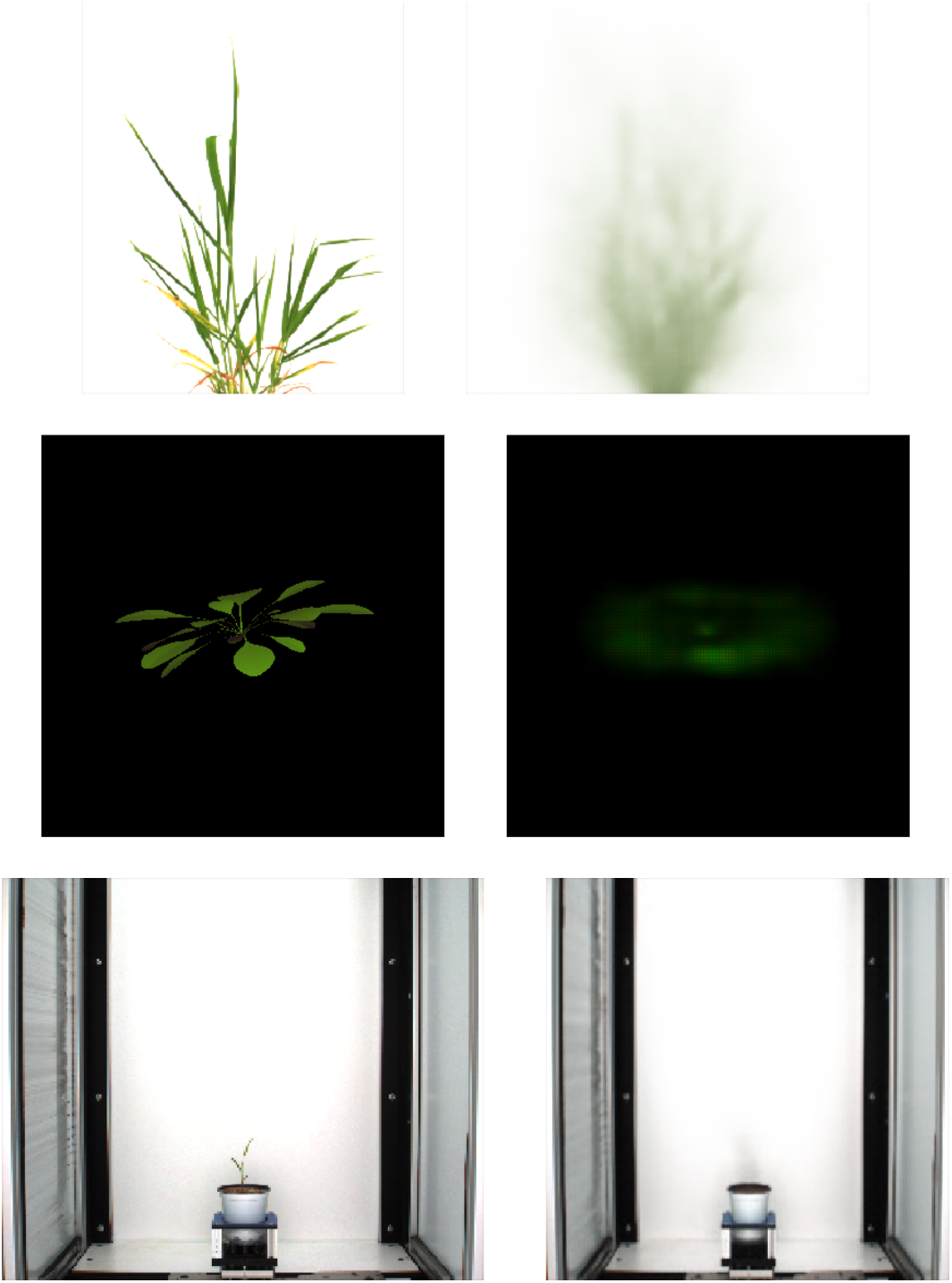
Real images from the *Setaria*, synthetic Arabidopsis, and sorghum datasets (left), and the same images predicted from their latent space encodings by a decoder network (right).

**Figure S4:**
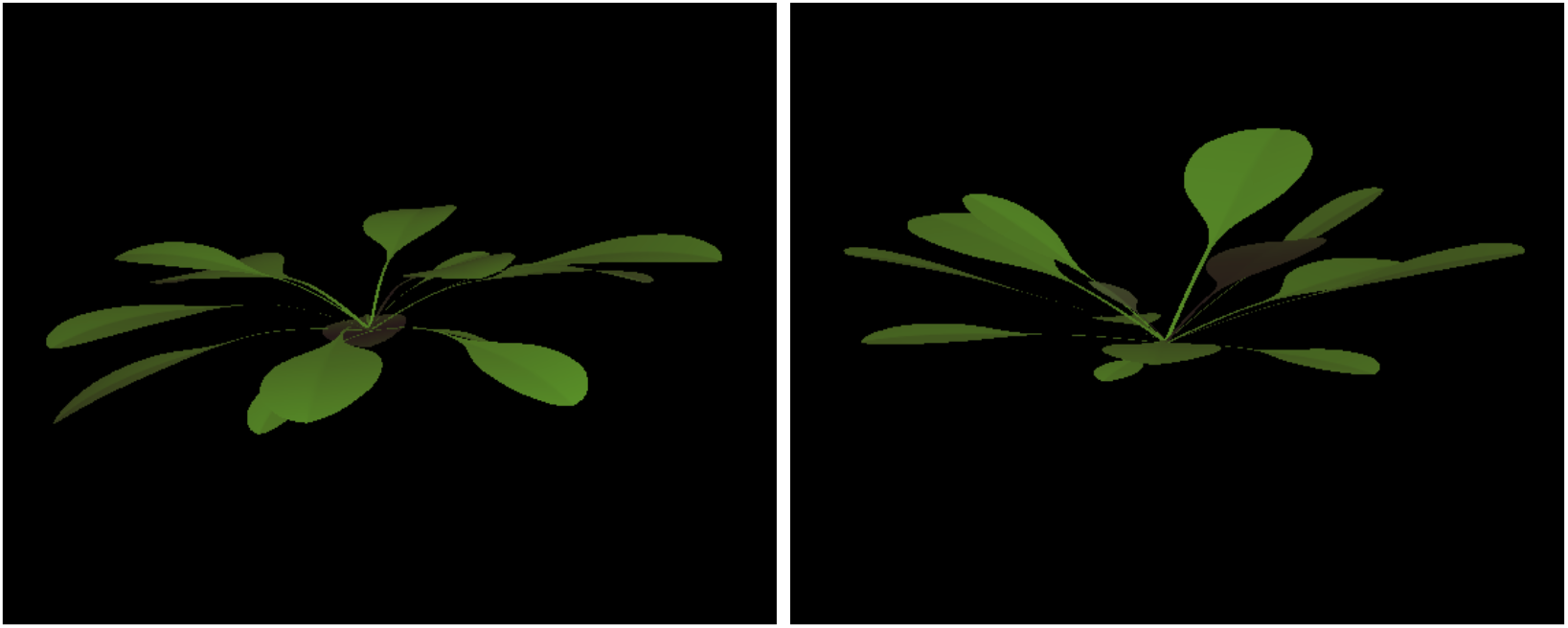
Untreated (left) and treated non-resistant (right) synthetic Arabidopsis plants at the final timepoint, showing differences in leaf elevation angle.

**Figure S5:**
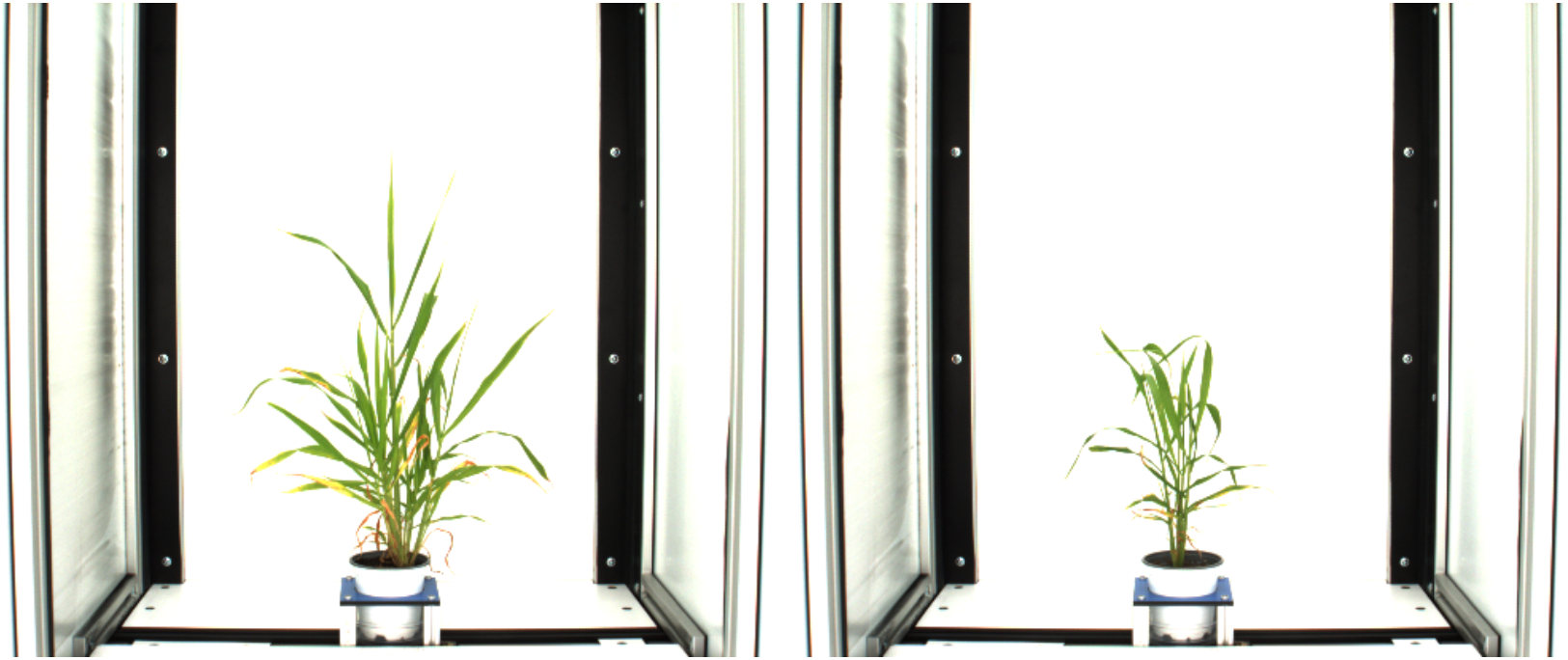
Well-watered (left) and water-limited (right) examples of a particular line from the *Setaria* RIL population [9]

**Figure S6:**
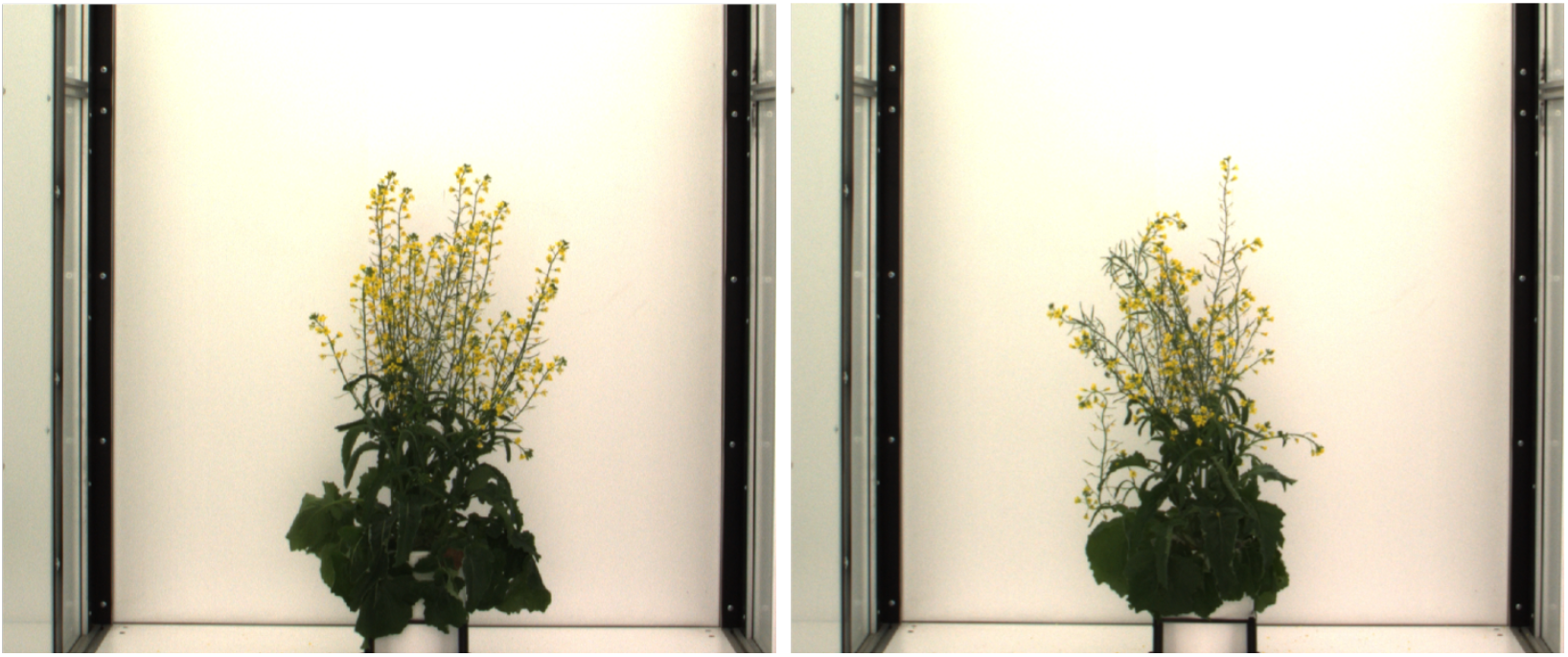
Well-watered (left) and water-limited (right) examples of a particular line from the *B. napus L*. NAM population

**Figure S7:**
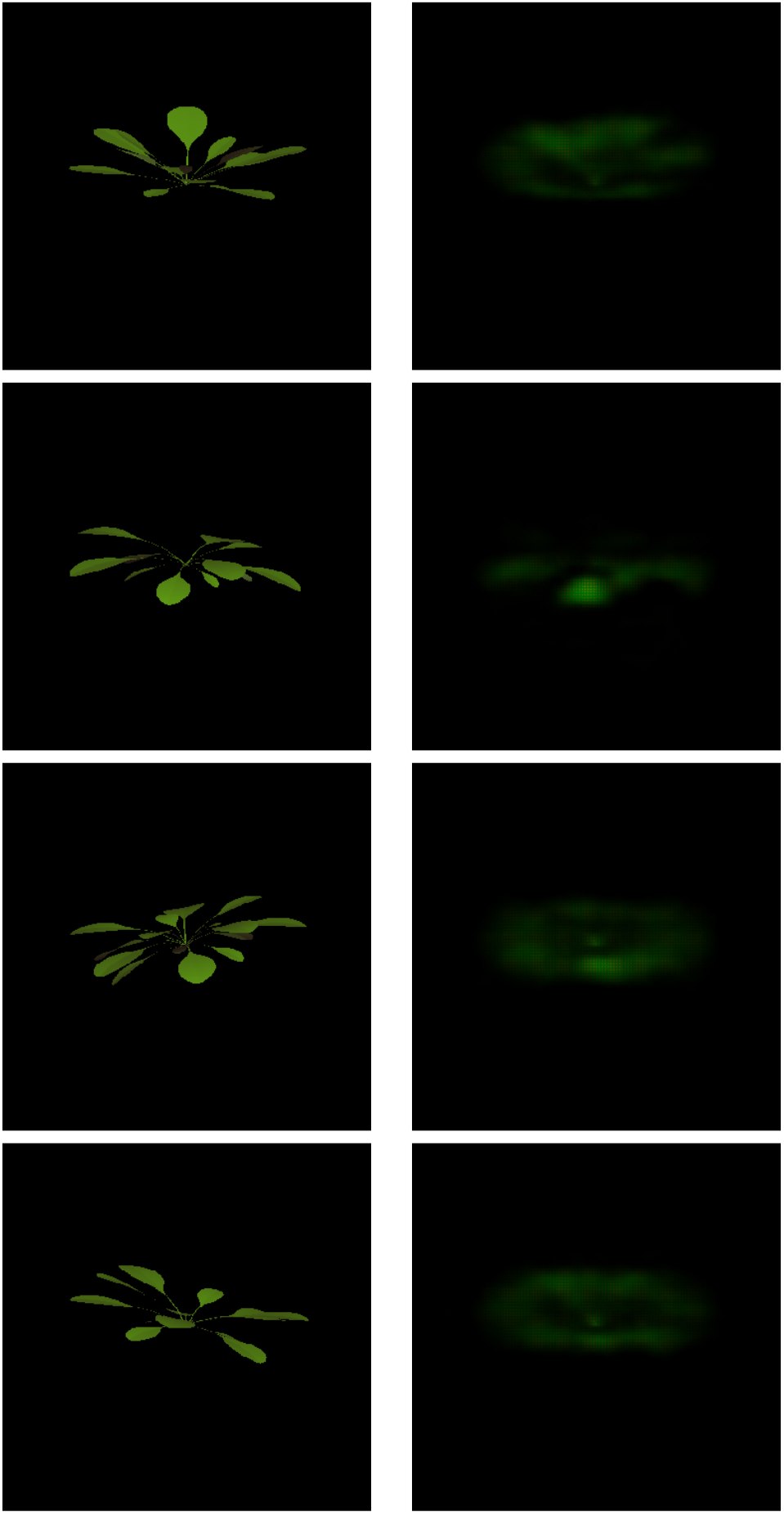
Additional examples of synthetic Arabidopsis rosettes (left) decoded from their latent space vectors (right).

**Figure S8:**
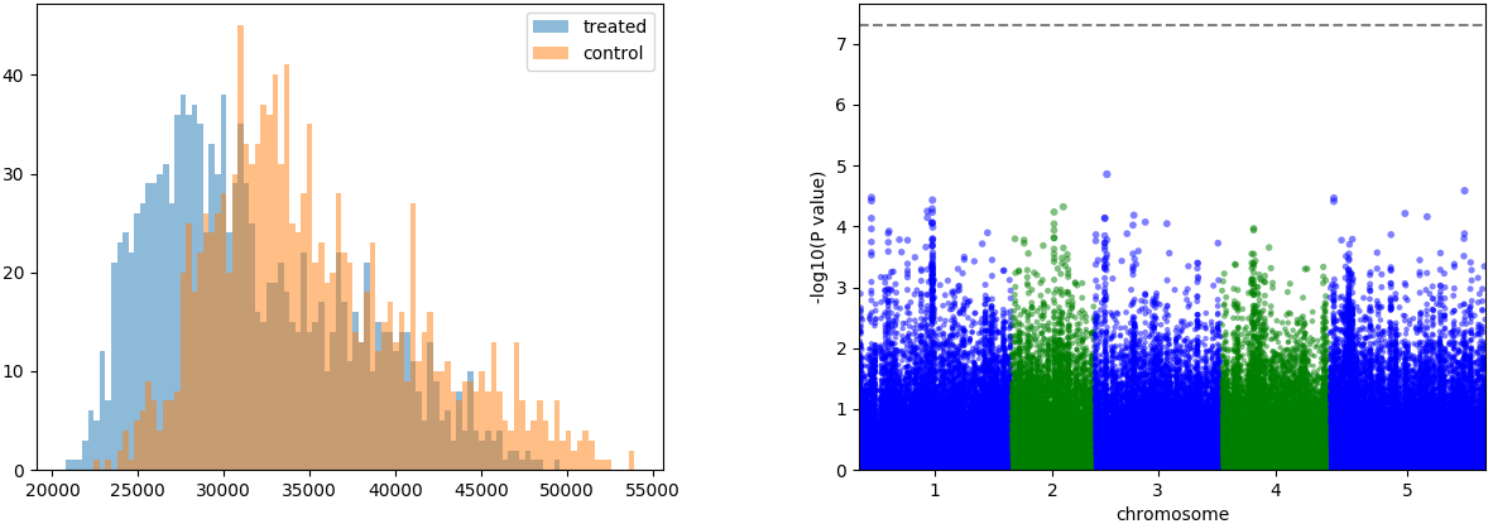
Ablation experiment using euclidean image distance between each pair of images in the sequence for the synthetic Arabidopsis dataset. The naive solution fails to recover the simulated resistance QTL on chromosome 1.

**Figure S9:**
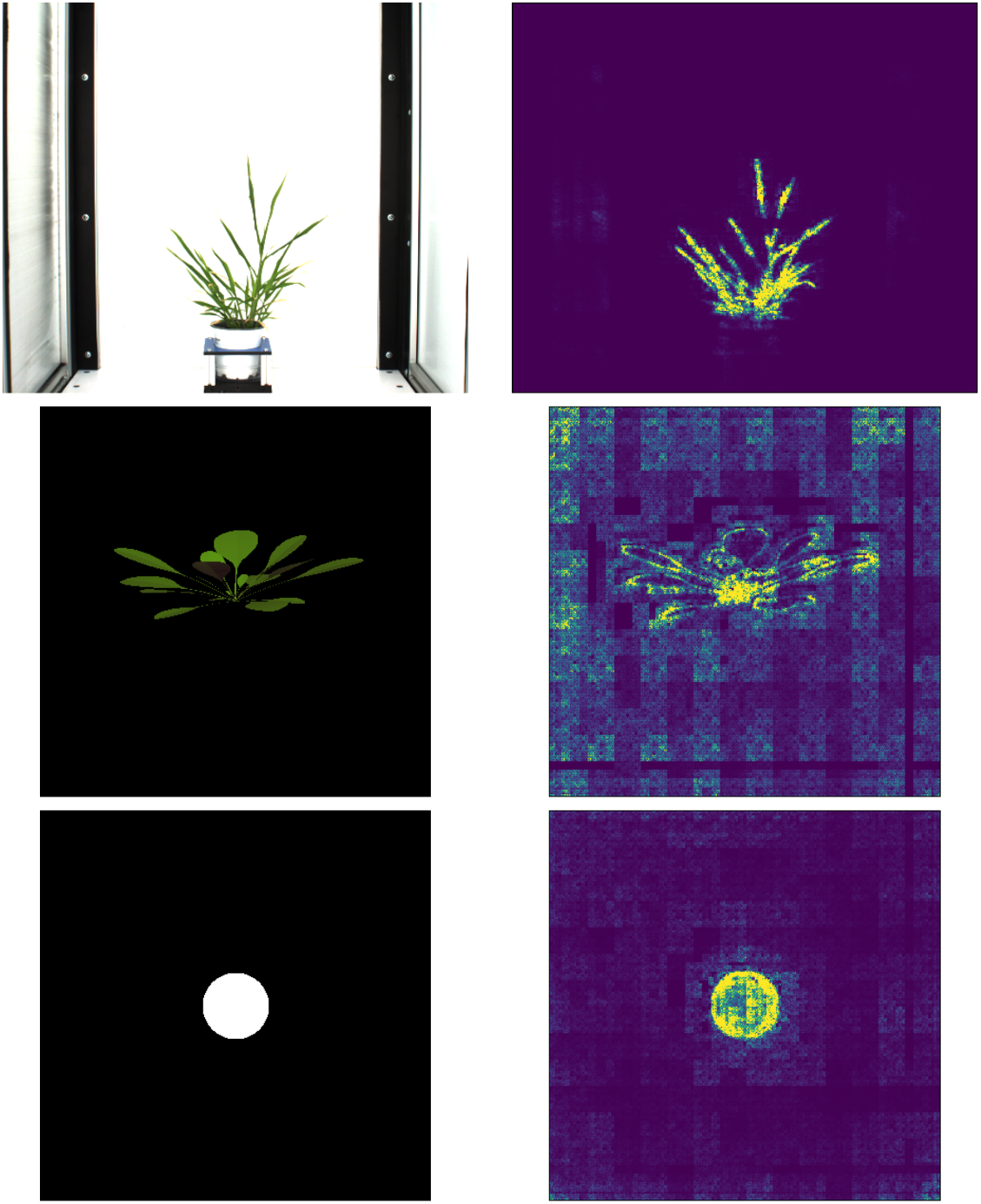
Example images from the *Setaria*, synthetic Arabidopsis, and synthetic circles datasets (left) and corresponding saliency maps generated using guided backpropagation (right). Intensity is higher for the pixels which have high saliency with respect to the L2 norm of the latent space embedding of the image. The *Setaria* image is from an experiment carried out without cropping to include the background in the saliency demonstration.

## References

[1] Virtual laboratory. http://www.algorithmicbotany.org/virtual_laboratory/. Accessed: 2017-08-01.

[2] Georgios Arvanitidis, Lars Kai Hansen, and Søren Hauberg. Latent Space Oddity: on the Curvature of Deep Generative Models. (2):1–15, 2017.

[3] Bernhard Biskup, Hanno Scharr, Ulrich Schurr, and Uwe Rascher. A stereo imaging system for measuring structural parameters of plant canopies. Plant, Cell & Environment, 30(10):1299–1308, 2007.

[4] Benjamin Brachi, Geoffrey P. Morris, and Justin O. Borevitz. Genome-wide association studies in plants: the missing heritability is in the field. Genome Biology, 12(10):232, Oct 2011.

[5] Malachy T. Campbell, Avi C. Knecht, Bettina Berger, Chris J. Brien, Dong Wang, and Harkamal Walia. Integrating Image-Based Phenomics and Association Analysis to Dissect the Genetic Architecture of Temporal Salinity Responses in Rice. Plant Physiology, 168(4):1476–1489, 2015.

[6] Tino Dornbusch, Séverine Lorrain, Dmitry Kuznetsov, Arnaud Fortier, Robin Liechti, Ioannis Xenarios, and Christian Fankhauser. Measuring the diurnal pattern of leaf hyponasty and growth in arabidopsis-a novel phenotyping approach using laser scanning. Functional Plant Biology, 39(11):860–869, 2012.

[7] Noah Fahlgren, Malia A. Gehan, and Ivan Baxter. Lights, camera, action: High-throughput plant phenotyping is ready for a close-up. Current Opinion in Plant Biology, 24:93–99, 2015.

[8] Max J. Feldman, Patrick Z. Ellsworth, Noah Fahlgren, Malia A. Gehan, Asaph B. Cousins, and Ivan Baxter. Components of Water Use Efficiency Have Unique Genetic Signatures in the Model C4 Grass Setaria. Plant Physiology, 178(2):699–715, 2018.

[9] Max J. Feldman, Rachel E. Paul, Darshi Banan, Jennifer F. Barrett, Jose Sebastian, Muh Ching Yee, Hui Jiang, Alexander E. Lipka, Thomas P. Brutnell, José R. Dinneny, Andrew D.B. Leakey, and Ivan Baxter. Time dependent genetic analysis links field and controlled environment phenotypes in the model C4 grass Setaria, volume 13. 2017.

[10] Robert T. Furbank and Mark Tester. Phenomics-technologies to relieve the phenotyping bottleneck. Trends in Plant Science, 16(12):635–644, 2011.

[11] Matthew W. Horton, Angela M. Hancock, Yu S. Huang, Christopher Toomajian, Susanna Atwell, Adam Auton, N. Wayan Muliyati, Alexander Platt, F. Gianluca Sperone, Bjarni J. Vilhjálmsson, Magnus Nordborg, Justin O. Borevitz, and Joy Bergelson. Genome-wide patterns of genetic variation in worldwide Arabidopsis thaliana accessions from the RegMap panel. Nature Genetics, 44(2):212–216, 2012.

[12] Andreas Kamilaris and Francesc X. Prenafeta-Boldú. Deep learning in agriculture: A survey. Computers and Electronics in Agriculture, 147(February):70–90, 2018.

[13] Diederik P. Kingma and Jimmy Lei Ba. Adam: a Method for Stochastic Optimization. International Conference on Learning Representations 2015, pages 1–15, 2015.

[14] Diederik P Kingma and Max Welling. Auto-encoding variational bayes. arXiv preprint arXiv:1312.6114, 2013.

[15] Samuli Laine. Feature-Based Metrics for Exploring the Latent Space of Generative Models. ICLR Workshop, 7:1–4, 2018.

[16] Yann LeCun, Yoshua Bengio, and Geoffrey Hinton. Deep learning. Nature, 521(7553):436–444, 2015.

[17] Dario Leister, Claudio Varotto, Paolo Pesaresi, Alexandra Niwergall, and Francesco Salamini. Large-scale evaluation of plant growth in Arabidopsis thaliana by non-invasive image analysis, 1999.

[18] Mao Li, Margaret H. Frank, Viktoriya Coneva, Washington Mio, Daniel H. Chitwood, and Christopher N. Topp. The persistent homology mathematical framework provides enhanced genotype-to-phenotype associations for plant morphology. Plant Physiology, 177(4):1382–1395, 2018.

[19] Christoph Lippert, Jennifer Listgarten, Ying Liu, Carl M. Kadie, Robert I. Davidson, and David Heckerman. FaST linear mixed models for genome-wide association studies. Nature Methods, 8(10):833–835, 2011.

[20] Guillaume Lobet, Iko T. Koevoets, Manuel Noll, Patrick E. Meyer, Pierre Tocquin, Loïc Pagès, and Claire Périlleux. Using a Structural Root System Model to Evaluate and Improve the Accuracy of Root Image Analysis Pipelines. Frontiers in Plant Science, 8(April):1–11, 2017.

[21] Hao Lu, Zhiguo Cao, Yang Xiao, Bohan Zhuang, and Chunhua Shen. TasselNet: Counting maize tassels in the wild via local counts regression network. Plant Methods, pages 1–14, 2017.

[22] Sharada Prasanna Mohanty, David P. Hughes, and Marcel Salathé. Using Deep Learning for Image-Based Plant Disease Detection. Frontiers in Plant Science, 7(September):1–7, 2016.

[23] Lars Mündermann, Yvette Erasmus, Brendan Lane, Enrico Coen, and Przemyslaw Prusinkiewicz. Quantitative modeling of Arabidopsis development. Plant Physiology, 139(2):960–968, 2005.

[24] E. H. Neilson, A. M. Edwards, C. K. Blomstedt, B. Berger, B. Lindberg Møller, and R. M. Gleadow. Utilization of a high-throughput shoot imaging system to examine the dynamic phenotypic responses of a c4 cereal crop plant to nitrogen and water deficiency over time. Journal of Experimental Botany, 66(7):1817–1832, 2015.

[25] Michael P. Pound, Jonathan A. Atkinson, Alexandra J. Townsend, Michael H. Wilson, Marcus Griffiths, Aaron S. Jackson, Adrian Bulat, Georgios Tzimiropoulos, Darren M. Wells, Erik H. Murchie, Tony P. Pridmore, and Andrew P. French. Deep machine learning provides state-of-the-art performance in image-based plant phenotyping. GigaScience, 6(10):1–10, 2017.

[26] Lufeng Qie, Guanqing Jia, Wenying Zhang, James Schnable, Zhonglin Shang, Wei Li, Binhui Liu, Mingzhe Li, Yang Chai, Hui Zhi, and Xianmin Diao. Mapping of Quantitative Trait Locus (QTLs) that contribute to germination and early seedling drought tolerance in the interspecific cross Setaria italica x Setaria viridis. PLoS ONE, 9(7):3–10, 2014.

[27] Bernardino Romera-Paredes and Philip Hilaire Sean Torr. Recurrent instance segmentation. In Bastian Leibe, Jiri Matas, Nicu Sebe, and Max Welling, editors, Computer Vision - ECCV 2016: 14th European Conference, Amsterdam, The Netherlands, October 11-14, 2016, Proceedings, Part VI, pages 312–329. Springer International Publishing, 2016.

[28] Asheesh Kumar Singh, Baskar Ganapathysubramanian, Soumik Sarkar, and Arti Singh. Deep Learning for Plant Stress Phenotyping: Trends and Future Perspectives. Trends in Plant Science, 23(10):883–898, 2018.

[29] J.T. Springenberg, A. Dosovitskiy, T. Brox, and M. Riedmiller. Striving for simplicity: The all convolutional net. In ICLR (workshop track), 2015.

[30] Sarah Taghavi Namin, Mohammad Esmaeilzadeh, Mohammad Najafi, Tim B. Brown, and Justin O. Borevitz. Deep Phenotyping: Deep Learning For Temporal Phenotype/Genotype Classification. bioRxiv, pages 1–29, 2017.

[31] Matthew A Turk and Alex P Pentland. Face recognition using eigenfaces. In Proceedings. 1991 IEEE Computer Society Conference on Computer Vision and Pattern Recognition, pages 586–591. IEEE, 1991.

[32] Jordan R. Ubbens, Mikolaj Cieslak, Przemyslaw Prusinkiewicz, and Ian Stavness. The use of plant models in deep learning: an application to leaf counting in rosette plants. Plant Methods, 14(1):6, 2018.

[33] Jordan R. Ubbens and Ian Stavness. Deep Plant Phenomics: A Deep Learning Platform for Complex Plant Phenotyping Tasks. Frontiers in Plant Science, 8(July), 2017.

[34] Kira M. Veley, Jeffrey C. Berry, Sarah J. Fentress, Daniel P. Schachtman, Ivan Baxter, and Rebecca Bart. High-throughput profiling and analysis of plant responses over time to abiotic stress. Plant Direct, 1(4):e00023, 2017.

[35] S.S. Wilks. Certain Generalizations in the Analysis of Variance. Biometrika, (24):471–494, 1932.

[36] Robail Yasrab, Jonathan A Atkinson, Darren M Wells, Andrew P French, Tony P Pridmore, and Michael P Pound. Rootnav 2.0: Deep learning for automatic navigation of complex plant root architectures. BioRxiv, page 709147, 2019.

